# *Chlamydomonas reinhardtii and Microbacterium forte sp. nov.,* a mutualistic association that favor sustainable hydrogen production

**DOI:** 10.1101/2023.05.03.539223

**Authors:** Neda Fakhimi, María Jesus Torres, Emilio Fernandez, Aurora Galván, Alexandra Dubini, David González-Ballester

## Abstract

A multispecies bacterial community including *Microbacterium forte* sp. nov., *Stenotrophomonas goyi* sp. nov., and *Bacillus cereus* greatly promoted sustained hydrogen production by the microalga *Chlamydomonas reinhardtii* when cocultivated in mannitol- and yeast extract-containing medium (up to 313 mL·L^-1^). Alga viability was also largely prolonged in the cocultures (>45 days) without any nutrient supplementation. Among the bacterial community, *Microbacterium forte* sp. nov. was the main responsible for the hydrogen production improvement. Nonetheless, the use of the entire bacterial community allowed a better growth of the alga during hydrogen production. *Chlamydomonas reinhardtii* and *Microbacterium forte* sp. nov. established a mutualistic association, based on the release of ammonium and acetic acid from the bacterium, while the alga provided sulfur-containing metabolites and complemented the bacterial auxotrophy for biotin and thiamine. This study uncovers the potential of the Chlamydomonas-bacteria consortia for durable and stable H_2_ production while allowing the simultaneous production of biomass.

## Introduction

Hydrogen (H_2_) is a very interesting clean fuel that do not release CO_2_ after combustion and that can be easily interconvertible with electricity, allowing the transport and storage of renewable energy (Abdin et al., 2020). The clean and sustainable H_2_ production (green H_2_) is currently part of the main energy policies of many governments (Kakoulaki et al., 2021). Among the different technologies to produce green H_2_, the biological production of H_2_ (bioH_2_) offers the possibility of culturing living organisms (bacteria, cyanobacteria, and algae) that liberate H_2._ This kind of technology, which is currently in TRL 3-4 stage, allows for H_2_ production at low temperatures (23-30°C) and atmospheric pressure, which is key to produce clean and sustainable energy at low costs (Lepage et al., 2021). BioH_2_ is particularly interesting when combined with other biological industrial applications such as wastewater treatment (Aydin et al., 2021) or any valorization of the biomass (e.g., biofertilizers, cosmetics, bioplastics, biofuels, etc.) (Fabris et al., 2020; Leong et al., 2021). Importantly, when using algae, the possibility of using light-driven energy and CO_2_ capture is also an advantage of this technology. Still, bioH_2_ is currently impeded by some technological (high cost, low TRL stage) and physiological barriers (low yield) that prevent its industrial application (Dębowski et al., 2020).

Heterotrophic bacteria link H_2_ production to their fermentative metabolism (dark H_2_) (Dahiya et al., 2021), while microalgae and cyanobacteria can produce H_2_ linked to their photosynthetic activity (photobioH_2_) (Ghirardi et al., 2009). In both cases, H_2_ production requires anoxic/hypoxic conditions since the hydrogenases/nitrogenases are oxygen sensitive. H_2_ production in microalgae have been widely studied in *Chlamydomonas reinhardtii* (Chlamydomonas throughout), a microalga able to grow photoautotrophycally or heterotrophically/mixotrophically in the presence of acetic acid (Chaiboonchoe et al., 2014). In this alga, there are two main pathways for H_2_ photoproduction, namely the PSII-dependent and PSII-independent pathways. The PSII-dependent uses electrons derived from water photolysis to produce H_2_, while the PSII-independent relies on the non-photochemical reduction of the plastoquinone pool (Fouchard et al., 2005; Hemschemeier et al., 2008). S deprivation have been widely employed to elicit sustainable PSII-dependent H_2_ production in Chlamydomonas, and many genetic engineered Chlamydomonas strains have been designed to improved H_2_ production during S deprivation (Dubini and Ghirardi, 2015). However, S deprivation is a stress condition that causes loss of cell viability, lowers biomass yield, and increases cultivation costs, which reduce the potential for industrial applications. On the other hand, H_2_ production in nutrient-replete media is also possible, mostly by controlling illumination and/or using acetate in the media (Degrenne et al., 2011; Scoma et al., 2014; Jurado-Oller et al., 2015). Assimilation of acetate under low light can elicit H_2_ production via the PSII-independent pathway (Jurado-Oller et al., 2015; González-Ballester et al., 2017). However, H_2_ production rates in nutrient-replete media are lower than during S-deprivation. In the last years, algae-bacteria cocultures have emerged as effective strategy to elicit H_2_ production in both S-depleted and nutrient-replete media (reviewed in (Fakhimi et al., 2020)).

In natural ecosystems, alga-bacteria associations can be complex and versatile, with many bidirectional metabolic collaborations consisting of the exchange of a variety of molecules (Hom et al., 2015; Ramanan et al., 2016; Yao et al., 2019). Bacteria can provide macronutrients like algal assimilable carbon (Cho et al., 2015) and nitrogen sources. The later can be provided by either fixing nitrogen (Kim et al., 2014) or mineralizing amino acids and peptides (Calatrava et al., 2018). Other nutrient exchanges from the bacteria to the algae include phytohormones (Teplitski and Rajamani, 2011), vitamins (Helliwell et al., 2011) and micronutrients (Amin et al., 2009). In return, the alga can supply O_2_ and fixed organic carbon sources (Cho et al., 2015b; Calatrava et al., 2018; Yao et al., 2019)

An evident advantage of coculturing heterotrophic bacteria with algae for H_2_ production, is that bacteria can efficiently remove O_2_ from the media, which is the most critical bottleneck associated with H_2_ photoproduction. However, other than O_2_ depletion, little information has been provided explaining the increased H_2_ production in algae-bacteria cocultures. In different Chlamydomonas-bacteria cocultures incubated with sugars, H_2_ photo-production was linked to the ability of the bacteria to produce acetic acid (Fakhimi et al., 2019a). Moreover, when H_2_ producing bacteria and Chlamydomonas are cocultured in sugar-containing media, a synergetic H_2_ production can be obtained by integrating the photobiological and fermentative production. This synergetic H_2_ production is based on acetic acid exchange (Fakhimi et al., 2019a).

This study describes a multispecies bacterial community composed of *Bacillus cereus* and two novel bacteria (*Microbacterium forte* sp. nov. and *Stenotrophomonas goyi* sp. nov.) capable of associating with Chlamydomonas enabling large sustained algal H_2_ production and long-term cell viability of the alga. The genome sequences and growth features of *M. forte* and *S. goyi* are discussed in the related publications (Fakhimi et al., 2023a; Fakhimi et al., 2023b). These two novel bacteria are unable to use sulfate as S source but require S-reduced forms such as cysteine and methionine to grow, while *M. forte* sp. nov. is additionally auxotroph for the vitamins biotin and thiamine (Fakhimi et al., 2023a; Fakhimi et al., 2023b). *Microbacterium forte* sp. nov. was the solely bacteria responsible of the increase in algal H_2_ production. The metabolic interactions established between *Microbacterium forte* sp. nov. and Chlamydomonas are analyzed and discussed.

## Materials and methods

### Algal and bacterial strains

*Chlamydomonas reinhardtii* strain 704 (cw15, arg7^+^, Nia1:Ars, mt^+^) (Loppes et al., 1999) was used in all the experiments. Chlamydomonas precultures was grown photomixotrophically on Tris Acetate Phosphate (TAP) medium (Harris, 2008) at 24°C under continuous Photosynthetic Photon Flux Density (PPFD) of 60-90 µmol photon·m^-2^·s^-1^. Three bacteria were identified in this work from a bacterial community. Individual members of this bacterial community were isolated by sequential rounds of plate streaking in Yeast Extract Mannitol (YEM) medium, until 3 different types of bacterial colonies were visually identified (*Supplemental Fig. 1*). Colonies were grown separately, and the subsequent isolated DNA was used for PCR-amplification of their partial RNA 16S sequences. After sequencing, the three isolated bacteria were identified as members of the genus *Microbacterium*, *Stenotrophomonas, and Bacillus*. Further PacBio whole genome sequencing and genome assembling by SNPsaurus LLC identified these three bacteria as *Microbacterium forte* sp. nov (Fakhimi et al., 2023a), *Stenotrophomonas goyi* sp. nov. (Fakhimi et al., 2023b), and *Bacillus cereus*. *Microbacterium forte* sp. nov. and *Stenotrophomonas goyi* sp. nov. have been deposited in the Spanish Type Culture Collection (CECT) with accession number CECT30765 and CECT30764, respectively. Genome sequences for *Microbacterium forte* sp. nov. and *Stenotrophomonas goyi* sp. nov. have been deposited in GenBank (NCBI) as CP116871 and SUB12685123. *M. forte* sp. nov. has been submitted for patenting (OEPM Submission P202330306)

All the bacterial precultures were grown on Yeast Extract Mannitol (YEM) medium or LB medium. The bacteria cultures were incubated at 25-28±1°C and under agitation (130 rpm).

### Coculturing alga and bacteria

Chlamydomonas precultures were grown to mid-log phase, harvested by centrifugation (3.000×g for 3 min) and resuspended in the corresponding media. Bacterial precultures grown until the Optical Density at 600 nm (OD_600_) reached 0.6-0.8 (logarithmic phase), then harvested by centrifugation (8.000×g for 3 min), washed twice and resuspended in the corresponding media to a final OD_600_ of 1.0. Alga and bacteria were mixed in the corresponding media with final chlorophyll concentration of 10 µg·mL^-1^ and bacterial OD of 0.05 or 0.1. The cocultures and their respective control monocultures were incubated in a growth chamber equipped with LED panels (AlgaeTron AG 230, Photon System Instruments) at 25°C and continuously agitated (180-200 rpm).

Media used in this work were: Minimal Medium (MM) (Harris, 2008); Tris Acetate Phosphate (TAP) (Harris, 2008); TAP supplemented with 56 mM of mannitol (TM); TAP supplemented with 0.8 g·L^-1^ of yeast extract (TY); TAP supplemented with 56 mM mannitol and 0.8 g·L^-1^ of yeast extract (TYM). In some experiments the mannitol and yeast extract concentrations of these media were lowered to 28 mM and 0.4 g·L^-1^, respectively, and the corresponding media denoted as TM_L_, TY_L_ and TYM_L_ (L standing for Low). Synthetic dairy wastewater was composed of TAP medium supplemented with 2.5 g·L^-1^ of tryptone and 2.5 g·L^-1^ of lactose.

Cultures used for growth test were performed in Erlenmeyer flasks (100 mL) containing 70 ml of the corresponding medium (termed as aerobic condition throughout). Cultures for H_2_ production were performed in hermetically sealed bioreactors (155 mL) containing 100 mL of culture media (termed as hypoxic condition throughout). Bioreactors for H_2_ production were opened under sterile conditions every 24 h (aeration) to release the H_2_ partial pressure and replace the gas composition in the headspace with atmospheric air (Jurado-Oller et al., 2015). The daily H_2_ production and O_2_ levels in the headspace were measured before aeration of the bioreactors. All cultures were continuously agitated under 50 μmol photons·m^-1^·s^-1^, unless otherwise is indicated.

### Measurement of alga and bacteria growth

Alga growth was assessed in terms of chlorophyll concentration (Wintermans and De Mots, 1965) and by counting cells number using a hemocytometer (Sysmex Microcellcounter F-500).

Bacteria growth in monocultures was estimated spectrophotometrically in term of OD_600_ evolution (DU 800, Beckman Coulter). In cocultures, bacterium cells were separated from alga cells by a customized Selective Centrifugal Sedimentation (SCS) approach (Torres et al., 2022). The goal of the technique consists in finding the centrifugation parameters that led to maximize algal cell sedimentation while minimize bacterial cell sedimentation. Thus, measuring the OD of the supernatant after centrifugation can provide an estimation of the bacterial growth in the cocultures. To do this, the % of precipitated cells of each monoculture were calculated at different forces (from 300 x g to 1000 x g) and times (30 s to 2 min) using the measured absorbances of individual monocultures before centrifugation (A_BC_) and after centrifugating the mixture of those cultures (A_AC_) (**Supplemental Table 1**). By using 500 x g for 1 min percentage of precipitated *M. forte* cells in cocultures was minimal (meaning that 95.6% of the *M. forte* cells remained in the supernatant). This condition was chosen as a good compromise for SCS and used to evaluate the contribution of the bacteria to the OD in cocultures (^SCS^OD_600_).

### Colony Forming Unit (CFU)

Colony Forming Unit (CFU) was quantified by making serial 10-fold dilutions of each culture, and 100 µl of each dilution was plated on LB plates. Plates were incubated for 3-4 days at 25°C. Later, the plates containing 50-100 colonies were selected, counted and further calculations were applied using corresponding dilution factors.

### Analyses of metabolites in gas and liquid phases

Gas chromatography (GC) (Agilent 7820A, Agilent Technologies) was used to detect H_2_ and O_2_ evolution in the headspace of the bioreactors. Gas samples from the headspaces (250 μl) were collected using a 1 mL Hamilton’s SampleLock™ syringe and manually injected into a Gas Chromatograph (GC) (Agilent 7820A, AgilentTechnologies). H_2_ and O_2_ were separated using a packed column (60/80 Molecular Sieve 5A, Ref. 13133-U, Supelco) at 75 °C and detected using a Thermal Conductivity Detector (TCD). Argon was used as the carrier gas.

Acetic acid, ethanol, lactic acid, formic acid, and mannitol analyses were performed by HPLC (Agilent series 1200, Agilent Technologies) using an ion-exchange column (Agilent Hi-Plex H, 300 × 7.7 mm, 6 μm I.D.) with isocratic elution using 5 mM H_2_SO_4_ at 50 °C. Samples (500 μL) were centrifuged, filtered (0.2 μm) and injected (20 μL) onto the HPLC system at a flow rate of 0.6 mL · min^-1^. Retention peaks were observed using UV (205 nm) and Refractive Index detectors.

## RESULTS

### Contaminated Chlamydomonas cultures enhanced H_2_ production in acetate-containing medium (TAP medium)

Preliminary experiments showed that the accidental bacterial contamination of Chlamydomonas TAP cultures enhanced H_2_ production (bacterial contamination derived from YEM medium). Supernatant from the contaminated Chlamydomonas cultures were preserved and used as inoculum for upcoming experiments. This Initial <u>B</u>acterial Inoculum will be referred as IBI throughout.

Chlamydomonas TAP cultures were inoculated with IBI in sealed bioreactors and used for H_2_ production experiments at different light intensities (12, 50 and 100 PPDF) (**Table 1**). As previously reported, Chlamydomonas control cultures produced more H_2_ at low light intensities (Jurado-Oller et al., 2015). In the IBI-Chlamydomonas cocultures, the H_2_ production was 1.4, 3.6 and 13.5 times higher than the respective monocultures incubated at 12, 50 and 100 PPDF, respectively. Similar results were obtained previously using cocultures of Chlamydomonas with different bacterial strains including *Pseudomonas putida, Pseudomonas stutzeri, E. coli* and *Rhizobium etli* where the highest improvements in H_2_ production, relative to Chlamydomonas monocultures, were obtained at high light intensities (Fakhimi et al., 2019b; Fakhimi et al., 2019a). No H_2_ production was observed in the solely bacterial IBI cultures, indicating that the bacterial community did not produce H_2_ under these conditions.

**Table 1:** H_2_ production (mL·L^-1^) in the TAP medium under different light intensities.

|  | 12 PPFD | 50 PPFD | 100 PPFD |
| --- | --- | --- | --- |
| **Chlamydomonas monocultures** | 15.2 ± 0.002 | 4.9 ± 0.02 | 0.4 ± 0.01 |
| **Chlamydomonas-IBI cocultures** | 21.6 ± 0.001 | 17.8 ± 0.02 | 5.4 ± 0.03 |
| **IBI monocultures** | 0 | 0 | 0 |

### The Chlamydomonas-IBI consortium can sustain H_2_ production and prolong the algal cell viability when incubated in the medium containing mannitol and yeast extract (TYM medium)

The H_2_ production capability of the Chlamydomonas-IBI cocultures was studied under different nutrient sources that are more suitable for the bacteria growth. TAP medium was either supplemented with mannitol (TM), yeast extract (TY) or both (TYM). H_2_ production, O_2_ evolution, growth, and acetic acid and mannitol uptake were monitored for 30 days (**Table 2** and **Fig. 1**).

**Figure 1.**
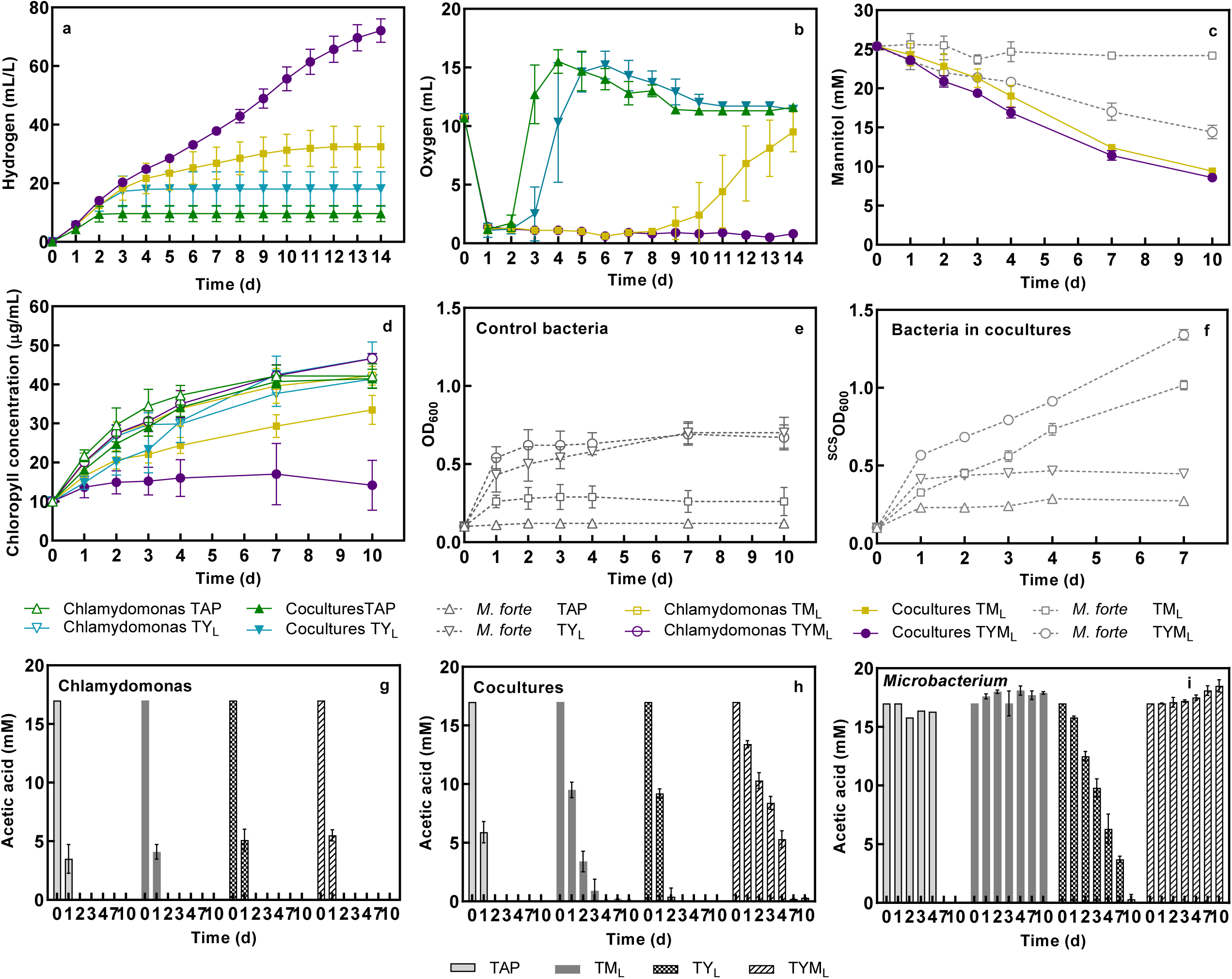
Comparison of Chlamydomonas-IBI cocultures in different media. H_2_ production (**a**), O_2_ evolution (**b**), chlorophyll concentration (**c**), bacterial OD_600_ (**d**), and mannitol (**e**) and acetic acid uptake (**f**) of Chlamydomonas-IBI (cocultures), and respective control cultures (algal monocultures and IBI cultures) incubated in TAP, TYM, TY or TM under 50 PPFD. For the sake of clarity, in panel (a) only cocultures are plotted. Mannitol and acetic acid uptake were not assayed in sole bacterial cultures. OD_600_ measurements were performed only over IBI cultures. Initial chlorophyll and bacterial OD_600_ were of 10 µg/mL and 0.05, respectively. Cultures were incubated in sealed bioreactors and aerated daily. H_2_ production in the headspaces was measured daily before aeration, and accumulative H_2_ evolution is plotted. Represented data are the average of three independent experiments.

**Table 2.**
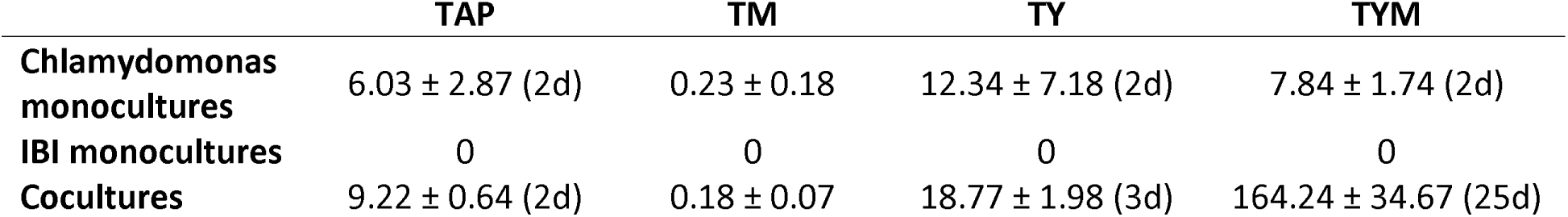
H production (mL·L^-1^) in different media. TAP was supplemented with 56 mM of mannitol (TM), or 0.8 g·L^-1^ of yeast extract (TY) or both (TYM). Numbers between parentheses indicated the length of the H_2_ production phase (days). Cultures were incubated under 50 PPFD.

|  | TAP | TM | TY | TYM |
| --- | --- | --- | --- | --- |
| **Chlamydomonas monocultures** | 6.03 ± 2.87 (2d) | 0.23 ± 0.18 | 12.34 ± 7.18 (2d) | 7.84 ± 1.74 (2d) |
| **IBI monocultures** | 0 | 0 | 0 | 0 |
| **Cocultures** | 9.22 ± 0.64 (2d) | 0.18 ± 0.07 | 18.77 ± 1.98 (3d) | 164.24 ± 34.67 (25d) |

All Chlamydomonas monocultures except TM monocultures produced H_2_ (6.03 to 12.34 mL·L^-1^) during the first two days, afterward no more H_2_ production was observed (**Table 2**). None of the IBI cultures produced H_2_, confirming that Chlamydomonas was the only H_2_ producer (**Table 2**). However, a very remarkable boost in H_2_ production was observed in cocultures incubated in TYM (164 mL·L^-1^; 21-fold total H_2_ production) (**Table 2** and **Fig. 1a**). A moderate improvement in H_2_ production was also observed in the TY cocultures (18.77 mL·L^-1^; 1.56-fold total H_2_ production). No H_2_ production was observed in the TM cocultures (**Table 2**). Moreover, H_2_ production in the TYM medium extended for 25 days, while in the rest of the cultures, the production was only sustained for 2-3 days. H_2_ production in the TYM cocultures experienced 3 phases. The first phase consisted in 2 days of relatively high H_2_ production rate (8 mL H_2_ ·L^-1^·d^-1^), afterwards 2 more days with very low H_2_ rate (< 1 mL·L^-1^·d^-1^), and then a third phase started from day 4 with increasing rates of H_2_ production to a maximum of 16.12 mL·L^-1^·d^-1^ on the 23^rd^ day (**Fig. 1a**). The TYM cocultures were the only Chlamydomonas-containing cultures able to sustain hypoxia for a long period (23 days) (**Fig. 1b**).

The growth of Chlamydomonas (by the mean of chlorophyll concentration) under H_2_ production condition was determined (**Fig. 1c**). In the TM medium, Chlamydomonas monocultures and cocultures turned yellow and died after 24h, indicating a likely toxic effect of the high mannitol concentration (56 mM) for the alga. Surprisingly, Chlamydomonas cells incubated in the TYM media did not suffer such a toxic effect. On the contrary, the chlorophyll content of the TYM monocultures and cocultures peaked at 76 and 59 μg·mL^-1^ on day 15, respectively, which was higher than in the standard TAP medium (48 μg·ml^-1^ for both monocultures and cocultures). These data suggest that the simultaneous supplementation with yeast extract and mannitol not only protected Chlamydomonas cells from the toxic effect of mannitol, but also provided a better growth condition relative to the standard TAP medium. Previously, it was shown that mannitol cannot be used as carbon source by Chlamydomonas (Chaiboonchoe et al., 2014; Fakhimi et al., 2019a). It may be possible that yeast extract is playing two roles simultaneously: a) compensating the potential osmotic stress caused by mannitol, and b) providing a N source for the alga through the activity of the extracellular L-amino acid oxidase (LAO1), which liberates ammonium and different α-keto-acids from small peptides and amino acids (Calatrava et al., 2019). This second role could also explain the higher growth observed in Chlamydomonas monocultures incubated in TY relative to TAP (**Fig. 1c**).

There was no difference in term of chlorophyll concentrations between the alga monocultures and cocultures incubated in TAP and TY, which reflect the neutral effect of the bacteria on Chlamydomonas growth in these media. However, in the TYM medium, less chlorophyll concentration was observed in the cocultures than in the monocultures. Still, a decline in chlorophyll concentration was observed in all the cultures in the 3^rd^ week, except in the TYM cocultures where the chlorophyll remained stable for 30 days (**Fig. 1c**), indicating a longer viability of the alga when cocultured with IBI in the TYM medium under hypoxia.

Solely bacterial cultures (IBI cultures) grew very efficiently in the presence of mannitol (TM and TYM media), showing a two-phase growth kinetic (**Fig. 1d**). During the first 5 days, the TYM and TY bacterial cultures grew similarly and faster than in TAP and TM. Afterwards, only TYM and TM showed a pronounced second growth phase reaching similar final ODs. The final growth of the cultures incubated in TAP and TY were also paired. Therefore, it appears that the presence of yeast extract promoted a rapid initial growth of the IBI cultures, although only the presence of mannitol allowed for sustained growth.

Despite mannitol not being assimilated by Chlamydomonas, continuous consumption of mannitol was observed in the TYM cocultures (**Fig. 1e**), indicating the ability of the bacteria community to uptake this nutrient. Acetic acid was quickly consumed in cocultures and algal monocultures (**Fig. 1f**). As previously reported, the slower the acetic acid is consumed the more sustained is hypoxia and H_2_ production in Chlamydomonas (Fakhimi et al., 2019b). Accordingly, the slightly slower consumption of acetic acid observed in the TY cocultures led to a bit more sustained H_2_ production relative to the TAP cocultures (3 vs 2 days) (**Table 2**). The TYM cocultures also showed a slower acetic acid uptake but a much more sustained H_2_ production phase (25 days). In this case, it is possible that the excretion of acetic acid from the bacteria through fermentation of mannitol supported sustained H_2_ production by the alga. Accumulation of acetic acid in the TYM cocultures was detectable only from day 18, although it is possible that experimentally undetectable levels of accumulated acetic acid were present from the 4^th^ day (**Fig. 1f**).

In another experimental set up, Chlamydomonas monocultures incubated in sealed bioreactors with TAP, TY and TYM were left for 16 days. During this period, a short (2 days) H_2_ production phase (< 12 mL·L^-1^) was obtained in all the cultures (**Fig. 2a**). Initial acetic acid was totally consumed after 16 days (**Fig. 2d**). On day 16, a set of cultures were inoculated with the IBI, and the rest remained as Chlamydomonas monocultures. A great and sustained H_2_ production was observed only in the IBI-inoculated cultures incubated in TYM (total of 313 mL H_2_ ·L^-1^ along 17 days) (**Fig. 2a**). O_2_ depletion was also observed only in the TYM-inoculated cultures (**Fig. 2b**). The TY and TAP cocultures did not produce extra H_2_ or deplete the O_2_. Acetic acid in the TYM medium was detectable 4 days after the IBI inoculation, which probably explains the algal H_2_ production observed in this medium (**Fig. 2d**). In all the cultures, but in the TYM cocultures, the chlorophyll content present on the 16^th^ day continuously dropped along the experiment (**Fig. 2c**). However, in the TYM cocultures the chlorophyll content remained constant (around 80 μg·mL^-1^) until the end of the experiment (day 30) and conserved a green color, confirming the previous observation that the IBI community can extend the viability of the Chlamydomonas cultures cultivated in the TYM medium under hypoxia.

**Figure 2.**
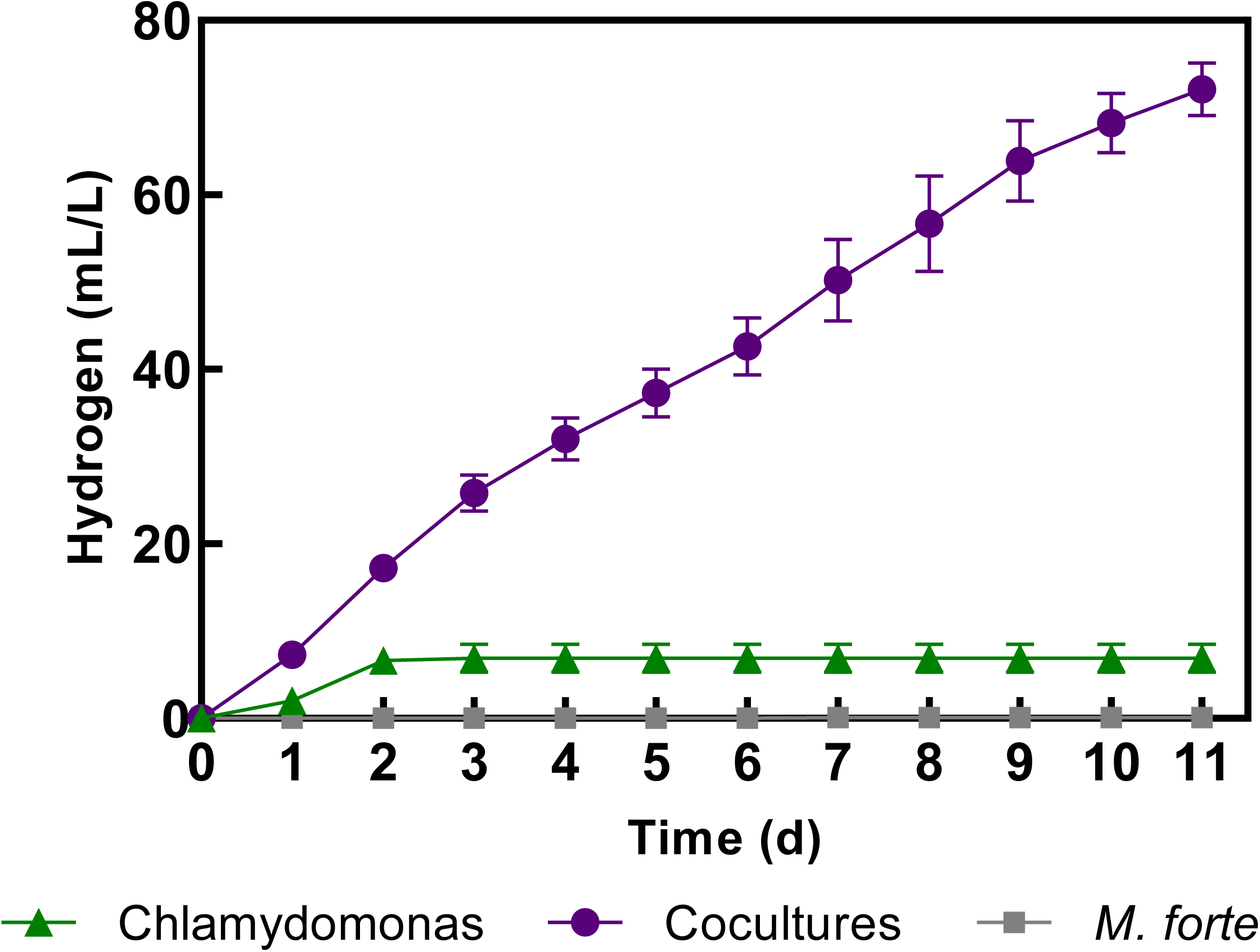
Effect of IBI inoculation on Chlamydomonas cultures. Chlamydomonas monocultures were incubated in TAP, TY and TYM media for 16 days (initial chlorophyll 10 µg·mL^-1^). On day 16^th^, a set of cultures were inoculated with the IBI (final OD_600_ of 0.05) (indicated with an arrow) and the rest were left as algal monocultures. H_2_ production (**a**), O_2_ evolution (**b**) during the 33 days and, chlorophyll concentration (**c**), and acetic acid levels (**d**) from day 15th are plotted. Cultures were incubated in sealed bioreactors and aerated daily. H_2_ production in the headspaces was measured daily before aeration, and accumulative H_2_ evolution is plotted. Data represent only one biological sample.

Overall, the Chlamydomonas-IBI cocultures incubated in TYM under hypoxia promoted sustained algal H_2_ production (up to 313 mL·L^-1^ for >16 days) (**Fig. 3a**) that was compatible with algal growth (up to 59 μg·mL^-1^) (**Fig. 1c**) and further extended Chlamydomonas cells viability for more than 30 days without any nutrient supplementation (**Fig. 1c and 2c**). This long viability of the Chlamydomonas cells was not observed in any other conditions. The extended viability could be linked to the acetic acid availability derived from the bacterial fermentation of mannitol (**Fig. 2d**), although the possibility that other nutrient exchanges (e.g., N or S sources), supporting Chlamydomonas survival cannot be discarded.

**Figure 3.**
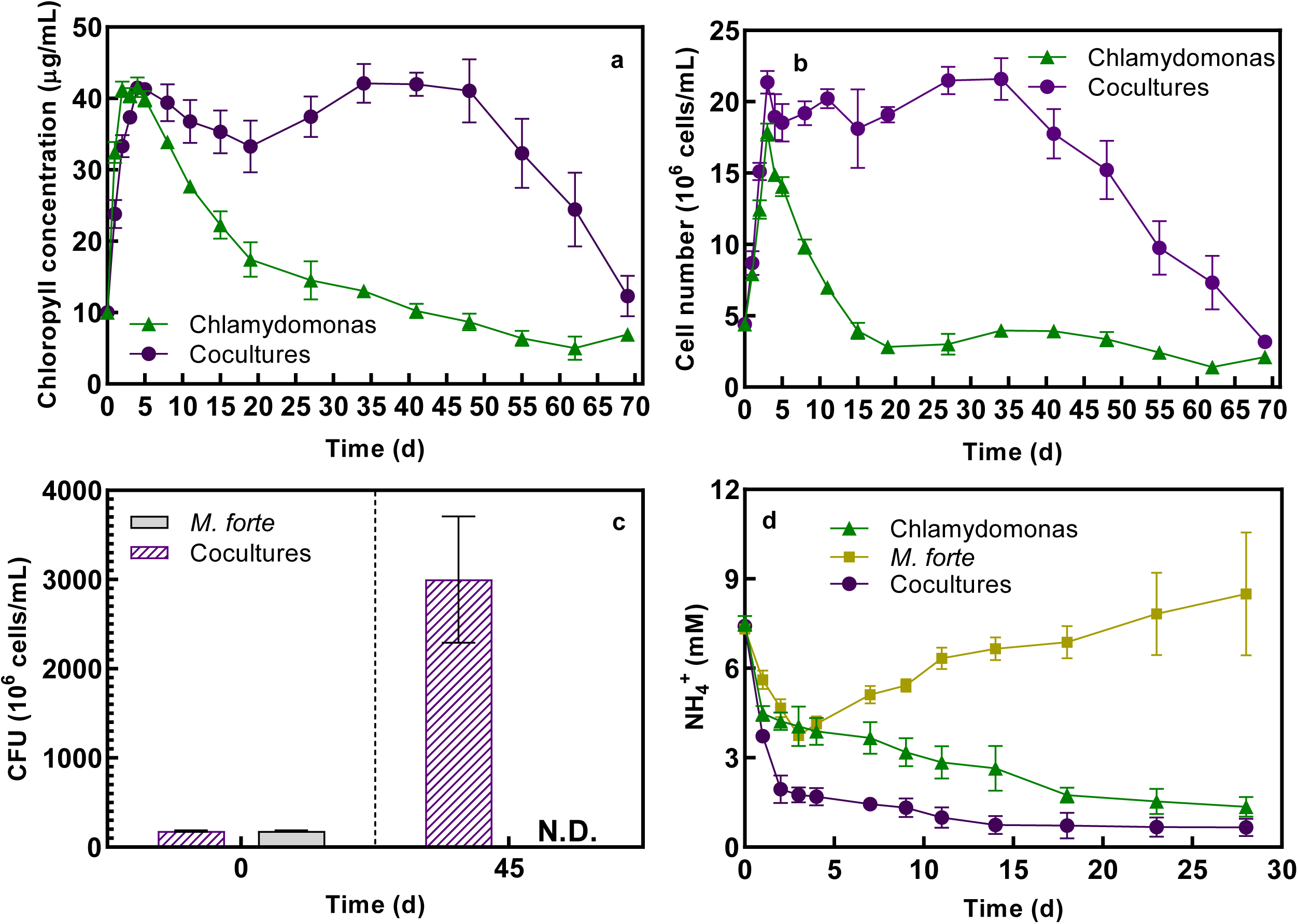
H_2_ production in cocultures of Chlamydomonas with *M. forte*., B*. cereus*., *S. goyi* or the original IBI community. H_2_ production (**a**), O_2_ evolution (**b**) of the different Chlamydomonas cocultures and respective monocultures, and mannitol (**c**) and acetic acid uptake (**d**) in bacterial monocultures. Chlamydomonas monocultures were incubated in the TYM medium, and on day 3, these cultures were inoculated with either, one, two or three different bacteria simultaneously (indicated by an arrow). Initial chlorophyll and bacterial OD_600_ were 10 µg/mL and 0.1, respectively. Cultures were incubated in sealed bioreactors and aerated daily. H_2_ production in the headspaces was measured daily before aeration, and accumulative H_2_ evolution is plotted. All cocultures are color-coded, while grey symbols refer to bacterial monocultures. Represented data are the average of three independent experiments.

To our knowledge, the H_2_ production detected in this consortium is the highest reported using non-genetically modified Chlamydomonas strains. Previously, the highest Chlamydomonas H_2_ production yields were obtained with the *pgr5* (850 mL·L^-1^; 9 days) (Steinbeck et al., 2015), *pgrl1* (437 mL·L^-1^; 5 days) (Tolleter et al., 2011), *stm6* (540 mL·L^-1^; 14 days) (Kruse et al., 2005), D1-*L159I-N230Y* (504 mL·L^-1^; 12 days) (Torzillo et al., 2009) and *Stm6Glc4L01* (361 mL·L^-1^; 8 days) (Oey et al., 2013) mutant lines in acetate-containing S-depleted medium (TAP-S).

The production of H_2_ observed in the Chlamydomonas-IBI cocultures is also the highest amount reported in any Chlamydomonas-bacteria cocultures (Fakhimi et al., 2020). Previously, the highest Chlamydomonas-bacteria H_2_ production was reported with *Bradyrhizobium japonicum* (170-141 mL·L^-1^; 14-16 days) (Wu et al., 2012; Xu et al., 2016) and *Pseudomonas* spp. (130 mL·L^-1^; 12 days) (Ban et al., 2018); all of them incubated under S-deficiency (TAP-S).

However, the use of mutant strains and/or S deficiency may have some biotechnological drawbacks. First, implementation of S deficiency (like any other nutrient limitation) implies a two-stage bioreactor process and often designing of synthetic media which are not the cheapest options, and greatly limit the utilization of inexpensive nutrient sources such as wastewaters. Moreover, H_2_ production in S-depleted media not only does not support growth but also declines the initial chlorophyll content and cell viability of the alga. Accordingly, no growth was reported during H_2_ production in the mutant strains described above. Moreover, all the mentioned above Chlamydomonas mutant strains have growth or/and photosynthetic deficiencies, which may further hinder the long-term sustainability of the cultures and the biomass production. Finally, genetically modified organisms could also raise some environmental concerns and face legal drawbacks that may further limit their use.

Importantly, the H_2_ production in the Chlamydomonas-IBI cocultures incubated in the TYM medium was concomitant with the growth of the alga and supported a long alga viability (more than in any other conditions) without any further nutrient supplementation.

### Identification of the members of the bacterial IBI community

To clarify the specific roles of each bacterium within the Chlamydomonas-IBI cocultures, 3 different bacteria present in this community were isolated and identified (**Supplemental Fig. 1**). Whole genome sequences were obtained and used to genetically identify and classified these 3 bacteria through the Type (Strain) Genome Server (TYGS) (Meier-Kolthoff and Göker, 2019). As a result, *Bacillus cereus* was identified together with two other novel species belonging to the genus of *Microbacterium* and *Stenotrophomonas*, which were named as *Microbacterium forte* sp. nov. (*M. forte*) (Fakhimi et al., 2023a) and *Stenotrophomonas goyi* sp. nov. (S. goyi) (Fakhimi et al., 2023b), respectively.

Interestingly, *Microbacterium sp*. and *Stenotrophomonas sp.* have been previously isolated from a contaminated Chlamydomonas culture with moderate enhanced H_2_ production in TAP-S (25-34 mL H_2_ ·L^-1^) (Li et al., 2013).

*Microbacterium sp., Bacillus* sp. and *Stenotrophomonas sp*. are among the most common inhabitants of the rhizosphere (Rosenblueth and Martínez-Romero, 2006; Marquez-Santacruz et al., 2010; Abadi et al., 2020). They are also recurrent bacteria frequently found within algal cultures (Burmølle et al., 2006; Li et al., 2013; Jones et al., 2016; Ji et al., 2019; Martelli et al., 2021; Mitra et al., 2021), which is probably reflecting frequent coexistence of these organisms in natural ecosystems.

### Deconvolution of the individual roles of each member of the IBI community in H_2_ production and extension of the alga viability

H_2_ production was evaluated in pairwise cocultures of Chlamydomonas with *M. forte* sp. nov., *S. goyi* sp. nov. or *B. cereus* in the TYM medium. Also, to test any potential multiple collaboration between the three bacteria, Chlamydomonas was cocultured with two-by-two bacteria combinations, with the three bacteria, as well as with the original IBI community (**Fig. 3**). Chlamydomonas monocultures were cultivated in the TYM medium for three days, afterwards, once the initial acetic acid content was totally consumed and the initial H_2_ production stopped, the different bacteria combinations were inoculated. Although all the cultures produced some H_2_, only the *M. forte*-containing cultures showed noticeable and sustained H_2_ production (**Fig. 3a**). H_2_ production among all the *M. forte*-containing cocultures was similar (35.8-47.1 mL·L^-1^) and sustained for at least 7 days. Moreover, although all the cocultures eventually reached hypoxia after bacteria inoculation, only the *M. forte*-containing cocultures were able to prolong hypoxia for several days (**Fig. 3b**). None of the bacteria monocultures produced H_2_, confirming that H_2_ was only produced by the alga. From these results, it was evident that only *M. forte* sp. nov. promoted sustained H_2_ production in Chlamydomonas, while *B. cereus* or *S. goyi* sp. nov. did not show a relevant role in H_2_ production.

The capability of the individual bacteria to uptake mannitol and acetic acid was also analyzed in the bacterial monocultures (**Fig. 3c** and **d**). Only *M. forte* and *B. cereus* were able to uptake some mannitol (**Fig. 3c**). *S. goyi* and *B. cereus* consumed acetic acid, especially the former which showed a very efficient acetic acid uptake (**Fig. 3d**). Importantly, the *M. forte* monocultures did not consume any acetic acid, but its excretion was observed, which probably derived from mannitol fermentation (**Fig. 3d**). This results also pointed out to *M. forte* as the only bacterium responsible for the acetic acid excretion observed in the Chlamydomonas-IBI cocultures (**Fig. 1f** and **2d**), which is also likely linked to the enhanced algal H_2_ production.

Once *M. forte* sp. nov. was identified as the only IBI member able to promote sustained H_2_ production in Chlamydomonas, the next question addressed was if this bacterium was also responsible for the prolonged viability of the Chlamydomonas cultures in the TYM medium. To answer this, Chlamydomonas and *M. forte* sp. nov. were cocultured in the TYM medium in not-sealed bioreactors (aerobic condition) for 70 days (**Fig. 4**). Due to the mannitol toxic effect on the alga previously encountered in the TM medium (**Fig. 1c**), the TYM mannitol concentration was reduced from 56 mM to 28 mM and referred as TYM_L_ (yeast extract was also lowered to 0.4 g·L^-1^ in the TYM_L_ medium). Initially, the Chlamydomonas monocultures and cocultures incubated in TYM_L_ grew similarly for 4-5 days (**Fig. 4a** and **4b**). After this initial growth, both the chlorophyll content and the alga cell number drastically decreased in the monocultures but not in the cocultures. The chlorophyll and cell number of the cocultures remained stable for more than 30 days (**Fig. 4a** and **4b**, and **Supplemental Fig. 2**), proving that *M. forte* sp. nov. can prolong the alga cell viability in the TYM_L_ medium under aerobic conditions.

**Figure 4.**
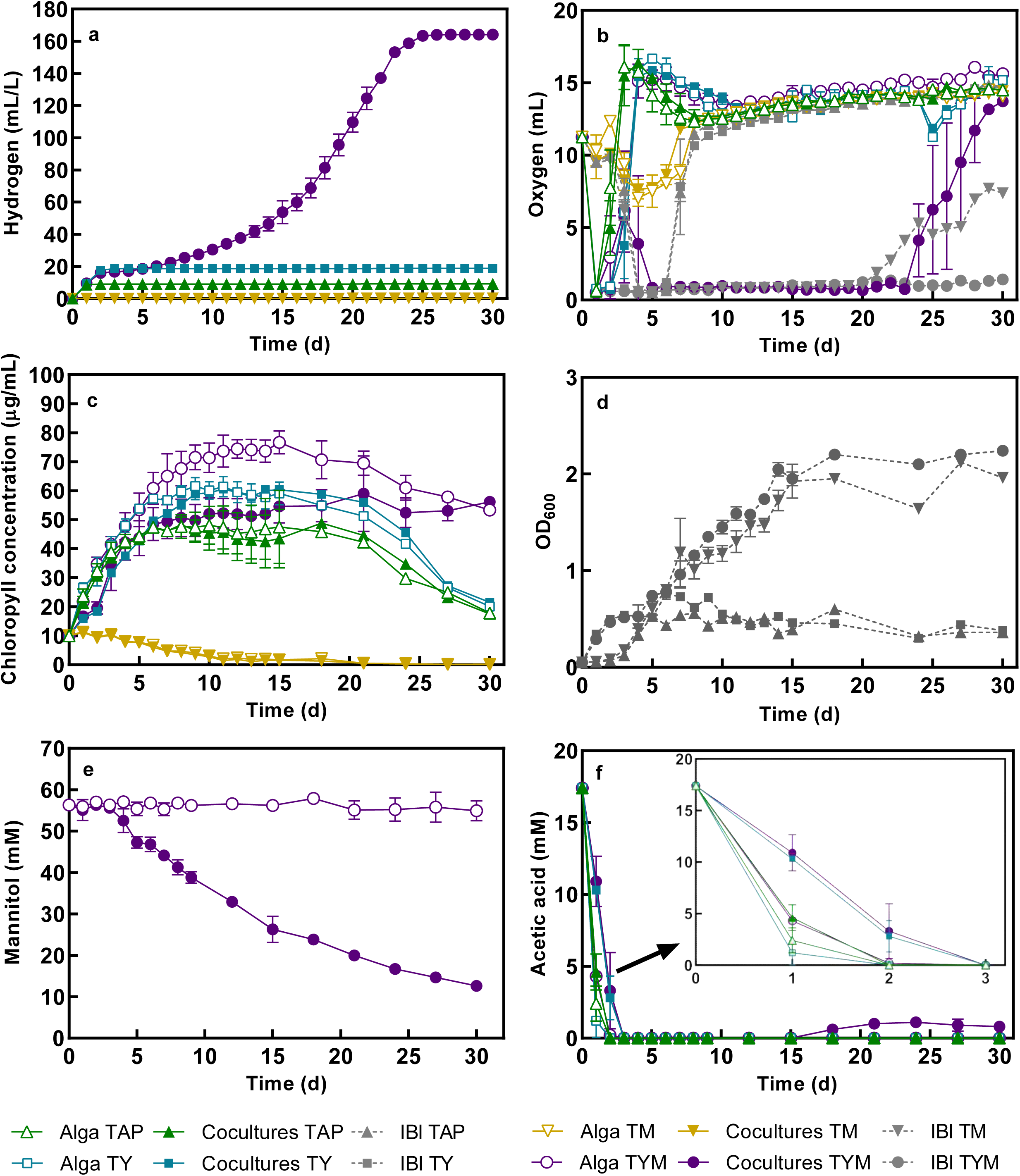
Long-term growth performance of Chlamydomonas-*M. forte* sp. nov. consortium under aerobic conditions. Chlorophyll content (**a**), Chlamydomonas cell number (**b**) and (**c**) bacterial Colony Forming Units (CFU), were determined in the Chlamydomonas-M. forte sp. nov. consortium and respective monocultures incubated in the TYM_L_ medium for 70 days. Ammonium released into the medium was also measured in cocultures under the same growth conditions for 28 days (**d**). CFU was measured on day 0 and 45. Initial chlorophyll and bacterial OD_600_ were 10 µg/mL and 0.1, respectively. Represented data are the average of three independent experiments. N.D., not detected.

As commented before, this extended algal cell viability is probably related with the capacity of *M. forte* sp. nov. to excrete acetic acid (Fig. 3d), which is used by Chlamydomonas as C source. But also, very important, the bacterial proteolytic activity led to the release of ammonium from amino acids/peptides, which provided the alga with a N source for a long period (Fig. 4d). Note that in *M. forte* sp. nov. monocultures, the ammonium accumulated in the medium after 30 days was above the initial ammonium concentration of the TYM medium (8 mM), indicating a great proteolytic capacity of this bacterium and a large ammonium availability in the medium broth.

Although the above results revealed a positive effect of *M. forte* sp. nov. on algal H_2_ production and alga viability, it remained unclear whether the bacterium also benefited from the alga (mutualism versus commensalism relationship). To clarify this situation, bacterial Colony Forming Units (CFU) were measured at the beginning of the experiment and on day 45 in both cocultures and bacteria control cultures. The CFU increased from 0.18×10^9^ to 3×10^9^ in cocultures, while no colonies appeared in the bacterial monocultures after 45 days (**Fig. 4c**). These results show that the alga and the bacterium established a mutualistic relationship where the growth of both microorganisms was benefited when incubated in TYM_L_. Similar results were obtained with *S. goyi* sp. nov. when cocultured with Chlamydomonas in TYM_L_ medium (Fakhimi et al., 2023b).

In sum, under aerobic conditions, Chlamydomonas and *M. forte* sp. nov. established a mutualistic relationship where the growth of both microorganisms was benefited. Similarly, in the Chlamydomonas-*Microbacterium paraoxydans* consortium used for phenol bioremediation, a significant increase in the growth rate of both microorganisms was found (Mora-Salguero et al., 2019). Yet, in another study, no alga growth promotion was observed in Chlamydomonas-*Microbacterium* sp. cocultures incubated in nutrient-repleted media (TAP) (Li et al., 2013).

Various strains of *Microbacterium* sp. are known to promote plant growth through indole-3-acetic acid and ammonia production, ACC deaminase activity (Singh and Singh, 2019) and by affecting S and N metabolism (Cordovez et al., 2018). However, the potential role and mechanisms of *Microbacterium* sp. as algal growth-promoting bacteria are unknown. In the Chlamydomonas-M. forte sp. nov. consortium incubated in TYM_L_, the extended algal viability observed under aerobiosis was likely supported by the acetic acid and ammonium secreted by the bacteria after metabolization of the mannitol and amino acids, respectively (**Fig. 3d, 5h and 5i**).

**Figure 5.**
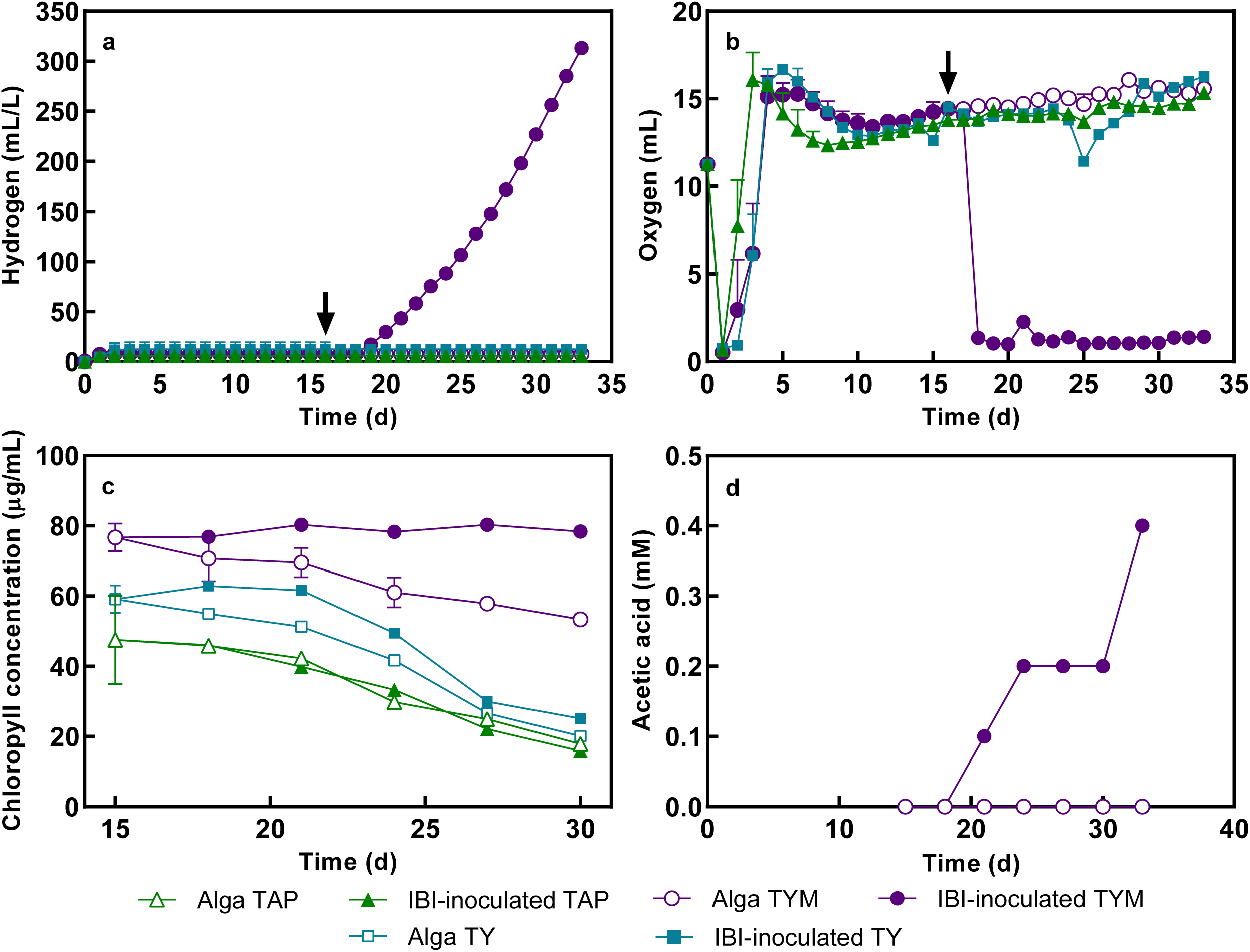
Comparison of Chlamydomonas-*M. forte* sp. nov. in different media. H_2_ production (**a**), O_2_ evolution (**b**), mannitol uptake (**c**) chlorophyll concentration (**d**), bacterial growth in monocultures (OD_600_) (**e**) and cocultures (OD_600_) (**f**), and acetic acid uptake (**f, h, i**) of Chlamydomonas-Microbacterium cocultures and respective monocultures. For the sake of clarity, in panel (a) only cocultures are plotted. Mannitol was not uptaken by Chlamydomonas monocultures (data not shown). Initial chlorophyll and bacterial OD_600_ were 10 µg/mL and 0.05, respectively. Cultures were aerated daily. H_2_ production in the headspaces was measured daily before aeration, and accumulative H_2_ evolution is plotted. Represented data are the average of three independent experiments.

On the other hand, in a related publication has been shown that *M. forte* sp. nov. is auxotroph for biotin and thiamine (Fakhimi et al., 2023a). Moreover, the bacterium can grow on different C sources although is unable to use sulfate as S source; its growth depends on S-reduced forms such as methionine or cysteine (Fakhimi et al., 2023a). Thereby, Chlamydomonas promoted the bacterium growth probably by alleviating the bacterial auxotrophy for biotin and thiamine and by providing S-reduced forms. Methionine and cystathionine are known S-containing metabolites secreted by Chlamydomonas (Vogel et al., 1978; Lorincz et al., 2010). In addition to these nutrients exchange, photosynthetic O_2_ probably also contributed to promote the bacterium growth in cocultures.

### Relevance of media composition during H_2_ production in the Chlamydomonas-M. forte sp. nov. consortium

H_2_ production in Chlamydomonas-*M. forte* sp. nov. cocultures and respective monocultures was assessed in sealed bioreactors (hypoxic condition) with different media: TAP, TM_L_, TY_L_ and TYM_L_ (**Fig. 5a**). As expected, H_2_ production was promoted and sustained in the TYM_L_ medium. Similarly, probably due to the decreased content of mannitol employed in the TM_L_ medium, and in turn the lower toxicity for Chlamydomonas, H_2_ was also produced in this medium, although less sustained and in less quantity compared with the production in the TYM_L_ medium. As previously observed, sustainability of the H_2_ production phase correlated very well with the presence of acetic acid in the medium (**Fig. 5h**) and the duration of the hypoxia phase (**Fig. 5b**).

Mannitol consumption in TM_L_ and TYM_L_ cocultures was similar (**Fig. 5c**). Mannitol uptake was also observed TYM_L_ bacterial monocultures, but not in TM_L_ bacterial monocultures (**Fig. 5c**). Accordingly, in the absence of yeast extract, *M. forte* sp. nov. monocultures did not grow efficiently on mannitol (TM_L_ medium) (**Fig 5e**). The growth of the bacterium in cocultures was estimated using a Selective Centrifugal Sedimentation (SCS) approach (Torres et al., 2022) (**Supplementary Table 1**). *M. forte* sp. nov. grew very efficiently on mannitol when co-cultivated with the alga, and independent of the addition of yeast extract (TYM_L_ and TM_L_ media) (**Fig. 5f**). From these results, it can be concluded that *M. forte* sp. nov. can grow efficiently on mannitol either when the medium is co-supplemented with yeast extract or when it is cocultured with Chlamydomonas (**Fig. 1c, 5e** and **5f**). Thereby, the alga must be providing some nutrients to the bacterium allowing the bacterial utilization of mannitol and growth in the absence of yeast extract. Indeed, mannitol uptake and bacterial growth was higher in the TM cocultures than in the TYM monocultures, implying a better performance of the bacterium when cocultured with the alga without yeast extract than when incubated as a monoculture in TYM. As commented before, Chlamydomonas is likely providing biotin, thiamine, cystathionine, and methionine that can support the bacterium growth on the medium without yeast extract (TM medium).

Moreover, *M. forte* sp. nov. monocultures were able to uptake acetic acid only in the TY_L_ medium (**Fig 5i**), which suggests that the bacterium only uptakes this nutrient if there is no other carbon source and yeast extract is available. Furthermore, the acetic acid was consumed faster in the TM_L_ than in the TYM_L_ cocultures (**Fig. 5h**). As commented before, slow acetic acid uptake rates benefit H_2_ production (Fakhimi et al., 2019b), which likely explains why the H_2_ production observed in TYM_L_ was higher than in TM_L_ (**Fig. 5a**). Thereby, the presence of yeast extract reduced the acetic acid uptake in the cocultures, which in turn benefited H_2_ production. Finally, the cocultures incubated in TYM_L_ showed higher accumulation of lactic acid and ethanol while less formic acid than the TM_L_ cocultures (**Supplemental Fig. 3**). These metabolic differences could also impact H_2_ production.

In sum, the presence of yeast extract in the media allowed the bacterial monocultures to uptake mannitol and grow. Nevertheless, in the absence of yeast extract, *M. forte* sp. nov. can also grow over mannitol if cocultivated with Chlamydomonas, which likely provides biotin, thiamine, cystathionine, and methionine. Still, the presence of yeast extract slowed down the uptake of acetate by the Chlamydomonas-*M. forte* sp. nov. cocultures, which is key to maximize H_2_ production.

Remarkably, no significant alga growth was observed in the TYM_L_ cocultures during H_2_ production condition (**Fig. 5d).** This fact greatly contrasts with the data obtained when Chlamydomonas was cocultivated with the IBI community under the same condition (hypoxia) (**Fig. 1c**) or with *M. forte* sp. nov. incubated under aerobic condition (**Fig. 4a**), where substantial alga growth was obtained. It is possible that other members of the IBI community (*Bacillus sp.* and/or *S. goyi*) counteract this negative impact of *M. forte* sp. nov. over the alga growth when cultivated under hypoxia.

### Potential of Chlamydomonas-*M. forte* sp. nov. consortium to produce H_2_ in a synthetic dairy wastewater

Due to the ability of *M. forte* sp. nov. to grow on sugars and peptides/amino acids, the potential of the consortium to produce H_2_ in synthetic dairy wastewaters was investigated. The consortium was cultivated in a synthetic dairy wastewater containing lactose and casein-derived tryptone. Sustained H_2_ production was obtained for at least 11 days, with a total production of 70 mL·L^-1^ (**Fig. 6**).

**Figure 6.**
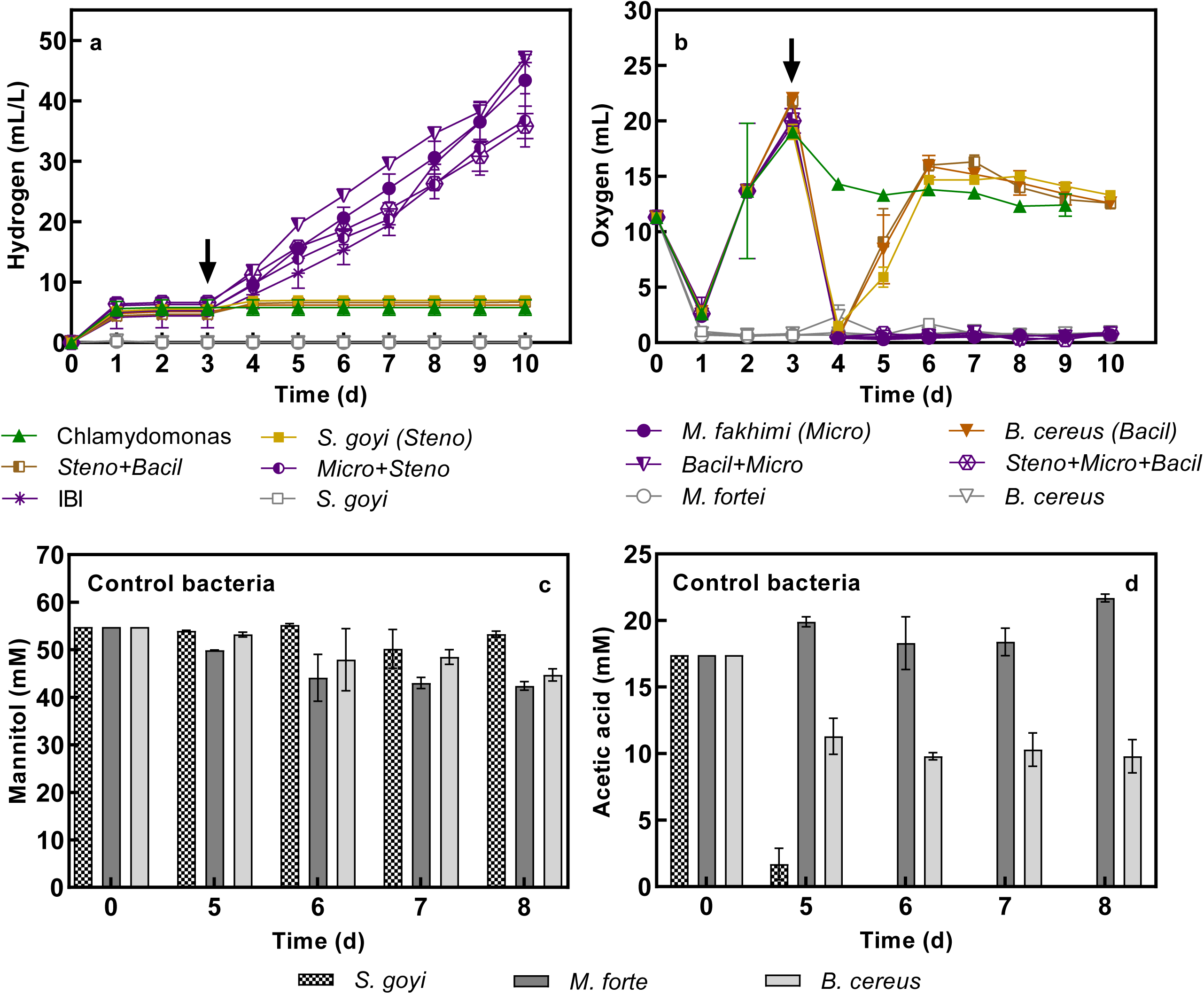
H_2_ production by Chlamydomonas-*M. forte* sp. nov. cocultures incubated in synthetic dairy wastewater. Synthetic dairy wastewater consisted of TAP medium supplemented with 2.5 g·L^-1^ of tryptone and 2.5 g·L^-1^ of lactose. Initial chlorophyll and bacterial OD_600_ were 10 µg/mL and 0.1, respectively. Alga and bacterium monocultures were used as controls. Cultures were aerated daily. H_2_ production in the headspaces was measured daily before aeration, and accumulative H_2_ evolution is plotted. Represented data are the average of three independent experiments.

This result highlights the potential biotechnological interest of Chlamydomonas-*M. forte* sp. nov. consortium.

### Suitability of the IBI- and *M. forte*-Chlamydomonas consortia for biotechnological purposes

Apart from the large sustained H_2_ production obtained with the IBI-Chlamydomonas consortium, one of the most interesting features of these consortium is the possibility of simultaneously obtaining H_2_ and algae biomass. Algae biomass is of biotechnological interest for the designing of new downstream process based on algae biomass valorization (Fabris et al., 2020; Leong et al., 2021), which can reduce the cost and thereby increase the economic viability of the industrial bioH_2_ production (Dębowski et al., 2020; Lepage et al., 2021). In this regard, BioH_2_ production and biomass valorization are particularly interesting when combined with wastewater treatment (Aydin et al., 2021). Some studies have shown the feasibility of producing bioH_2_ in wastewater environment, although the H_2_ production rate quickly and considerably declined (Hwang et al., 2018; Shetty et al., 2019). In the Chlamydomonas-IBI and Chlamydomonas-*M. forte* cocultures sustainable H_2_ production is possible if the media is enriched with sugar and proteins/peptides. Dairy wastewater could be an ideal broth to assay concomitant bioH_2_ production, bioremediation, and biomass valorization with these consortia, although further research with real dairy wastewater is needed to confirm this potential.

When using different Chlamydomonas-bacteria consortia, various parameters have been associated with the interruption of algal H_2_ production: a) rapid death of bacteria (Lakatos et al., 2014); b) acetic acid depletion (Lakatos et al., 2014; Fakhimi et al., 2019b); and c) loss of alga cells viability due to low pH and/or excessive accumulation of bacterial fermentative products, especially of acetic acid (Fakhimi et al., 2019a). For example, in Chlamydomonas-*E. coli* consortia incubated in glucose, there is a rapid acetic acid accumulation and pH decrease that provoke the alga death. On the other hand, in Chlamydomonas cocultures with *P. putida* or *R. etli* incubated with glucose and mannitol, respectively, acetic acid accumulation in the media was low and in turn the alga produced low H_2_ rates (Fakhimi et al., 2019a).

All these drawbacks were successfully bypassed in the Chlamydomonas-IBI and -*M. forte* sp. nov. consortia incubated in TYM: a) the bacterial growth was promoted in cocultures relative to bacterial monocultures, at least for *M. forte* and *S. goyi* (**Fig. 4c** and **5e,f**) (Fakhimi et al., 2023b); b) out of the three members of the IBI community only *S. goyi* showed intense acetic acid uptake when incubated in TYM, whereas *M. forte* excreted acetic acid (**Fig. 3d**), which allowed sustained algal H_2_ production (**Fig. 1a** and **5a**); and c) the mannitol fermentation, especially by *M. forte* sp. nov., was fast enough to provide sufficient acetic acid allowing sustainable H_2_ production (∼5.5 mL·L^-1^·d^-1^), but slow enough to avoid acidification of the medium and impair the viability of the alga. In fact, algal viability was largely promoted in the Chlamydomonas-IBI and Chlamydomonas-*M. forte* sp. nov. cocultures under hypoxic and aerobic conditions, respectively.

On the other hand, it is intriguing that the Chlamydomonas-*M. forte* cocultures, unlike the Chlamydomonas-IBI cocultures, did not promoted the alga growth when incubated under hypoxia (**Fig. 5d**). Hence, the presence of *S. goyi* sp. nov. and *B. cereus* could be crucial to obtain a growth equilibrium that benefited the alga growth and the H_2_ production in TYM_L_ medium under hypoxia.

The potential metabolic complementation that these four microorganisms could established in TYM_L_ medium could be complex and interdependent. Interestingly, *S. goyi* sp. nov. and *M. forte* sp. nov. are unable to use sulfate as S source and they show a dependency on S-reduced sources such as cysteine and methionine (Fakhimi et al., 2023a; Fakhimi et al., 2023b). Hence, the survival of *M. forte* sp. nov. and *S. goyi* sp. nov. in natural ecosystems could be highly dependent on the surrounding microbial communities. In this multispecies association, the alga could provide S-reduced sources in the form of cystathionine and methionine to both *S. goyi* sp. nov. and *M. forte*. Moreover, Chlamydomonas, *S. goyi* and *B. cereus* could be alleviating the auxotrophy for biotin and thiamine of *M. forte* sp. nov. (the genomes of *S. goyi* sp. nov. and *B. cereus* have entire biosynthetic pathways for biotin and thiamine, whereas the biosynthesis of these vitamins is well documented in Chlamydomonas (Aaronson et al., 1977; Croft et al., 2007)). On the other hand, the bacteria could provide CO_2_ and acetic acid to the alga through their fermentative activity. Acetic acid is mostly provided by *M. forte* sp. nov. Moreover, *S. goyi* sp. nov. and *M. forte* have great proteolytic capacity and can grow over amino acids/peptides as only C and N sources (Fakhimi et al., 2023b; Fakhimi et al., 2023a). Thereby, they both could also provide the alga with ammonium derived from the mineralization of the amino acids/peptides.

In any case, this multispecies association was mutually beneficial and guaranties the viability of the consortium by preventing an excessive bacterial growth, which could be one of the main drawbacks when algae-bacteria cocultures are used for biotechnological applications. Further research is needed to better resolve the relationships of these four microorganisms when incubated together.

## Conclusion

The consortium composed of the alga Chlamydomonas, *M. forte* sp. nov., *S. goyi* sp. nov. and B. cereus can concomitantly sustain H_2_ and biomass production when incubated in mannitol and yeast extract. *M. forte* sp. nov. is the sole bacterium responsible to promote H_2_ production in Chlamydomonas, although *S. goyi* sp. nov. and B. cereus could be important to generate alga biomass and extend the alga viability under hypoxia. *M. forte* sp. nov. and *S. goyi* sp. nov. can be mutualistically associated with the alga Chlamydomonas. This consortium could be of biotechnological interest for H_2_ and biomass production using wastewaters. The potential biotechnological application and the precise metabolic complementation of this consortium need to be further investigated under different growth conditions.

---

**Supplemental Figure 1.** Visual characterization of the IBI community under the microscope (**a**), and the isolated colonies of *Stenotrophomonas goyi* sp. nov. (**b**), *Bacillus cereus* (**c**), and *Microbacterium forte* sp. nov. (**d**).

**Supplemental Figure 2.** Chlamydomonas monocultures and Chlamydomonas-M. forte cocultures incubated in TYM along 60 days.

**Supplementary Figure 3.** Lactic acid (**a**, **b**, **c**, **d**), formic acid (**e**, **f**, **g**, **h**) and ethanol (**i**, **j**, **k**, **l**) concentrations found in the media of Chlamydomonas-*M. forte* sp. nov. cocultures and respective monocultures incubated in TAP, TM_L_, TY_L_, TYM_L_. Growth conditions are as in Fig 5. Represented data are the average of three independent experiments.

**Supplemental Table 1. Centrifugation forces and times used to determine the Selective Centrifugal Sedimentation (SCS) approach for Chlamydomonas and *M. forte*.**

Absorbance for Chlamydomonas (750nm) and *M. forte* monocultures (600nm) were measured as monocultures before centrifugation (A_BC_). Both monocultures were mixed (1:1) and measured again after centrifugation (A_AC_) at a specific centrifugation force and time. Dilutions factors were applied. Error % = 100 – [(A_AC_· 100)/A_BC_]. This value indicates the percentage of the error obtained in estimating the absorbance of Chlamydomonas or *M. forte* cultures in the mixed cultures compared to their initial monocultures (before mixing). The optimized centrifugation force (x g) and time (min) used for the Chlamydomonas and *M. forte* consortium is highlighted in bold. Data represent the means from at least two different biological samples and assayed with technical duplicates.

## Author Contributions

Writing—original draft preparation, D.G.-B and N.F..; writing—review & editing, N.F and A.D.; Bacteria isolation, N.F.; design and execution of experiments, N.F., M.J.T.; in silico analysis, D.G.-B.; results analysis and interpretation, N.F., D.G.-B.; supervision, A.D. and D.G.-B.; project administration, A.G., E.F., A.D. and D.G.-B.; funding acquisition, A.G., E.F., A.D. and D.G.-B. All authors have read and agreed to the published version of the manuscript.

## Acknowledgments

The authors acknowledge Dr. Gregorio Galvéz Valdivieso for his unvaluable contribution to this research: perfection sometimes kills new discoveries.

