## Supplementary figures and images for "*Chlamydomonas reinhardtii and Microbacterium forte sp. nov.,* a mutualistic association that favor sustainable hydrogen production"

### Supplemental Figure 2

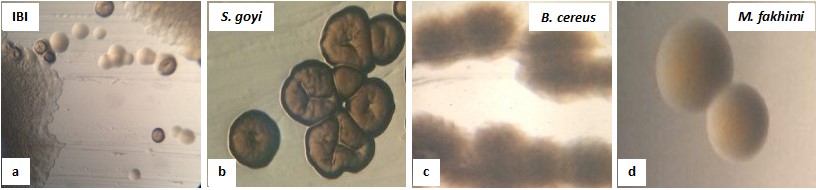

### Supplemental Figure 2

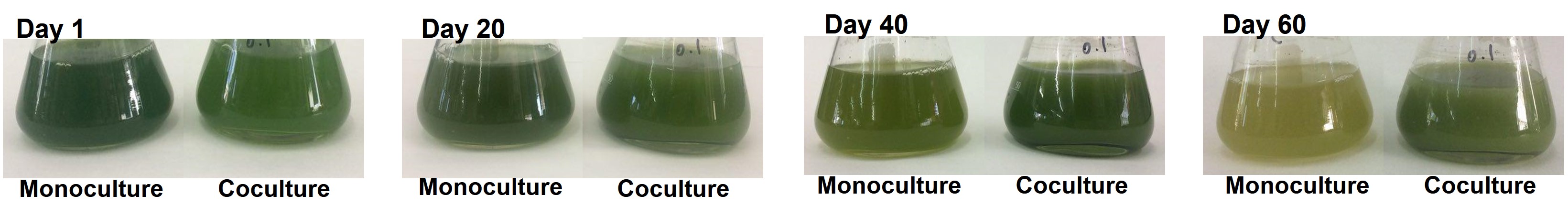

### Supplemental Figure 3

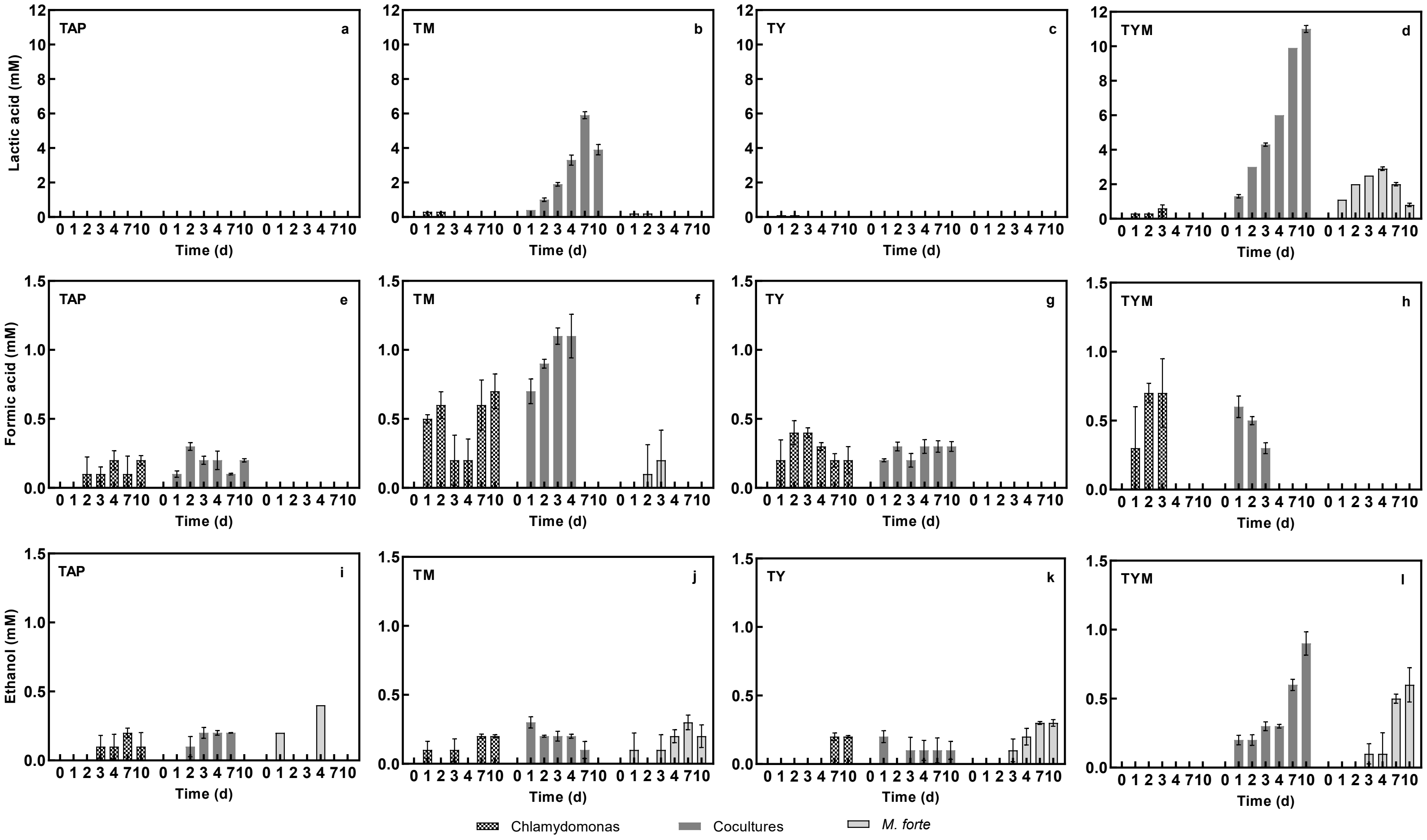
