## Supplemental Table for "*Chlamydomonas reinhardtii and Microbacterium forte sp. nov.,* a mutualistic association that favor sustainable hydrogen production"

|  |  | Chlamydomonas | | | *Microbacterium sp.* | | |
| --- | --- | --- | --- | --- | --- | --- | --- |
| Force (g) | Time (s) | A_BC_ | A_AC_ | Error % | A_BC_ | A_AC_ | Error % |
| 1000 | 60 | 0,92 | 1,30 ± 0,15 | -40,58 ± 19,13 | 0,82 | 0,436 ± 0,18 | 46,83 ± 22,07 |
| 1000 | 30 | 0,92 | 1,20 ± 0,13 | -29,87 ± 14,08 | 0,82 | 0,646 ± 0,08 | 21,22 ± 10 |
| 500 | 120 | 0,92 | 1,24 ± 0,15 | -33,77 ± 15,92 | 0,82 | 0,532 ± 0,13 | 35,12 ± 16,56 |
| **500** | **60** | 0,92 | 0,92 ± 0,00 | **0,49 ± 0,23** | 0,82 | 0,856 ± 0,02 | **-4,39 ± 2,07** |
| 500 | 30 | 0,92 | 0,87 ± 0,02 | 5,36 ± 2,52 | 0,82 | 1,12 ± 0,14 | -36,77 ± 17,33 |
| 400 | 120 | 0,92 | 1,13 ± 0,10 | -22,73 ± 10,71 | 0,82 | 0,71 ± 0,05 | 13,35 ± 6,29 |
| 400 | 60 | 0,92 | 0,87 ± 0,03 | 5,84 ± 2,75 | 0,82 | 0,85 ± 0,02 | -4,21 ± 1,98 |
| 300 | 120 | 0,92 | 1,04 ± 0,06 | -12,66 ± 5,97 | 0,82 | 0,78 ± 0,02 | 4,57 ± 2,15 |
| 300 | 60 | 0,92 | 0,85 ± 0,04 | 8,12 ± 3,83 | 0,82 | 1,17 ± 0,17 | -42,99 ± 20,26 |

**Supplemental Table 1. Centrifugation forces and times used to determine the Selective Centrifugal Sedimentation (SCS) approach for Chlamydomonas and *M. forte.***

Absorbance for Chlamydomonas (750nm) and *M. forte* monocultures (600nm) were measured as monocultures before centrifugation (A_BC_). Both monocultures were mixed (1:1) and measured again after centrifugation (A_AC_) at a specific centrifugation force and time. Dilutions factors were applied. Error % = 100 – [(A_AC_ · 100)/A_BC_]. This value indicates the percentage of the error obtained in estimating the absorbance of Chlamydomonas or *M. forte* cultures in the mixed cultures compared to their initial monocultures (before mixing). The optimized centrifugation force (x *g*) and time (min) used for the Chlamydomonas and *M. forte* consortium is highlighted in bold. Data represent the means from at least two different biological samples and assayed with technical duplicates.
